# Human Hippocampal Theta Oscillations Organise Distance to Goal Coding

**DOI:** 10.1101/2024.12.12.628182

**Authors:** Zimo Huang, James A Bisby, Neil Burgess, Daniel Bush

**Affiliations:** Department of Neuroscience, Physiology and Pharmacology, UCL, London, WC1E 6BT, United Kingdom; Division of Psychiatry, UCL, London, W1T 7BN, United Kingdom; UCL Queen Square Institute of Neurology, UCL, London WC1N 3BG, United Kingdom; UCL Institute of Cognitive Neuroscience, UCL, London, WC1N 3AZ, United Kingdom; Wellcome Centre for Human Neuroimaging, UCL, London, WC1N 3AR, United Kingdom

## Abstract

The rodent hippocampal local field potential is dominated by 6-12 Hz theta oscillations during active behaviour that are strongly implicated in spatial coding and memory function across species. Invasive electrophysiology in both rodents and humans has shown increases in hippocampal theta power immediately before the onset of translational movement that persists throughout subsequent motion, and the magnitude of this increase correlates with the distance subsequently travelled. Using non-invasive magnetoencephalography (MEG) and an abstract navigation task, we observed increased theta power during both spatial planning and subsequent navigation. Importantly, theta power in the right hippocampus covaried with subsequent path distance during planning, only when participants were aware of the distance to their goal. During subsequent navigation, hippocampal theta power decreased dynamically as participants approached the goal, only when they were aware of how far they still needed to travel. In addition, theta phase during navigation modulated 70-140 Hz fast gamma amplitude in the entorhinal cortex while traversing novel paths; and 30-70 Hz slow gamma amplitude in the right hippocampus while traversing previously experienced paths. In both cases, theta-gamma phase-amplitude coupling increased with proximity to the goal during navigation, consistent with the hypothesis that sequences of upcoming locations were represented by gamma bursts occurring at successive theta phases. In sum, these findings are consistent with the proposed role of hippocampal theta oscillation in flexible planning and goal-directed spatial navigation across mammalian species.

## Introduction

The mammalian hippocampus plays a central role in episodic memory and spatial cognition – the crucial ability to know where we are and where we are going (Scoville and Milner, 1957, Rosenbaum et al., 2000, Maguire et al., 2006, O’Keefe and Nadel, 1978). In rodents, the hippocampal local field potential (LFP) is dominated by 6-12 Hz theta oscillations during active engagement with the environment (Vanderwolf, 1969, O’Keefe and Nadel, 1978). Both the frequency (Sławińska and Kasicki, 1998, Jeewajee et al., 2008, Rivas et al., 1996) and power (Vanderwolf, 1969, McFarland et al., 1975) of theta oscillations covary with running speed, suggesting a potential role in encoding self-motion information. An increased prevalence of hippocampal theta oscillations has also been observed in human intracranial recordings during navigation in virtual (Kahana et al., 1999) and real-world (Aghajan et al., 2017, Bohbot et al., 2017) environments. Intriguingly, electrophysiology studies in both rodents and humans have shown that this increase in theta power is often greatest immediately prior to the onset of movement (Bush et al., 2017, Kaplan et al., 2012, Schmidt-Hieber and Häusser, 2013, Whishaw and Vanderwolf, 1973, Bland et al., 2006, Fuhrmann et al., 2015). Moreover, it has been demonstrated that movement-related theta power may correlate with distance to the goal both prior to (Bush et al., 2017, Vass et al., 2016) and during (Liu et al., 2023) virtual navigation. These findings support the hypothesis that hippocampal theta oscillations might encode navigationally relevant variables across mammalian species.

In humans, theta oscillations over the frontal midline are also implicated in cognitive control and working memory function (Cavanagh and Frank, 2014). For example, frontal theta power has been shown to increase with working memory load (Jensen and Tesche, 2002), and it has been suggested that theta-gamma ‘phase-amplitude coupling’ may support the neural coding of sequential information (Lisman and Idiart, 1995, Lisman and Jensen, 2013). According to this model, individual items are encoded by transient bursts of gamma band activity that are sequentially embedded in successive phases of each theta cycle (Lisman and Jensen, 2013, Jensen and Colgin, 2007). As the sequence of items being encoded becomes longer, gamma power becomes more widely distributed across the theta cycle, leading to a decrease in theta-gamma phase-amplitude coupling (Heusser et al., 2016, Daume et al., 2024). In the rodent hippocampus, theta oscillations have been shown to modulate gamma power in two distinct bands: a slower band, generated locally and implicated in memory retrieval; and a faster band, originating in the entorhinal cortex and implicated in the encoding of new information (Colgin et al., 2009). In addition, hippocampal place cells fire at progressively earlier phases of each theta cycle as their firing fields are traversed (Burgess et al., 1994, O’Keefe and Recce, 1993). This can generate ‘theta sweeps’ of place cell activity in each oscillatory cycle that correspond to spatial trajectories ahead of the animal (Wikenheiser and Redish, 2015, Burgess et al., 1994, Skaggs et al., 1996, Johnson and Redish, 2007). Under the framework described above, theta sweeps that encode longer trajectories would correspond to a wider distribution of gamma power across the theta cycle, and therefore lower phase-amplitude coupling.

Here, we asked healthy participants undergoing MEG to perform a self-paced navigation task in which they must learn the spatial layout of an abstract map of images while planning routes to specific goal images within that map. First, we demonstrate that theta power during spatial planning correlates with distance to the goal, only during trials in which a direct path to the goal is known. In addition, we observe a dynamic reduction in theta power during subsequent navigation as participants approach the goal. Source localisation revealed that this distance-to-goal coding originates from the bilateral hippocampus. Second, we demonstrate a dynamic increase in theta-gamma phase-amplitude coupling during navigation as participants approach the goal. Importantly, we find that theta phase modulates fast gamma amplitude in the entorhinal cortex during learning, when participants traverse new routes for the first time; and slow gamma amplitude in the hippocampus during subsequent retrieval, when participants retrace previously experienced paths. In sum, these findings support the hypothesis that theta oscillations scaffold the encoding of spatial trajectories across mammalian species, providing novel insight into the neural mechanisms of goal-directed navigation.

## Methods

### Participants

Twenty-seven healthy participants were recruited to participate in this experiment. Ethical approval was granted by the local research ethics committee at University College London. All participants gave written informed consent and were compensated for taking part. Two participants were excluded due to issues with the behavioural task, one was excluded for poor performance, and one for excessive MEG signal artefact. Of the remaining twenty-three participants, sixteen were male and eighteen were right-handed, with a mean ± SD age of 24.1 ± 4.89 years (range 18-35 years).

### Experimental Design

In the abstract navigation task, participants moved across a 4 x 4 grid comprised of 16 images to reach a specific goal location in each trial (Fig. 1A, B). Participants were instructed to navigate to the goal location using the shortest possible path - moving up, down, left, or right in each step using a button box. Importantly, participants never saw the whole map, which remained fixed across the entire experiment: at any time, only the four locations that were immediately north, south, east, and west of their current location were shown on screen (except at the edge or corners of the map, where fewer locations were available). As such, participants had to learn the map layout during navigation in each trial. The correspondence between map locations and visual stimuli was randomised for each participant.

**Figure 1:**
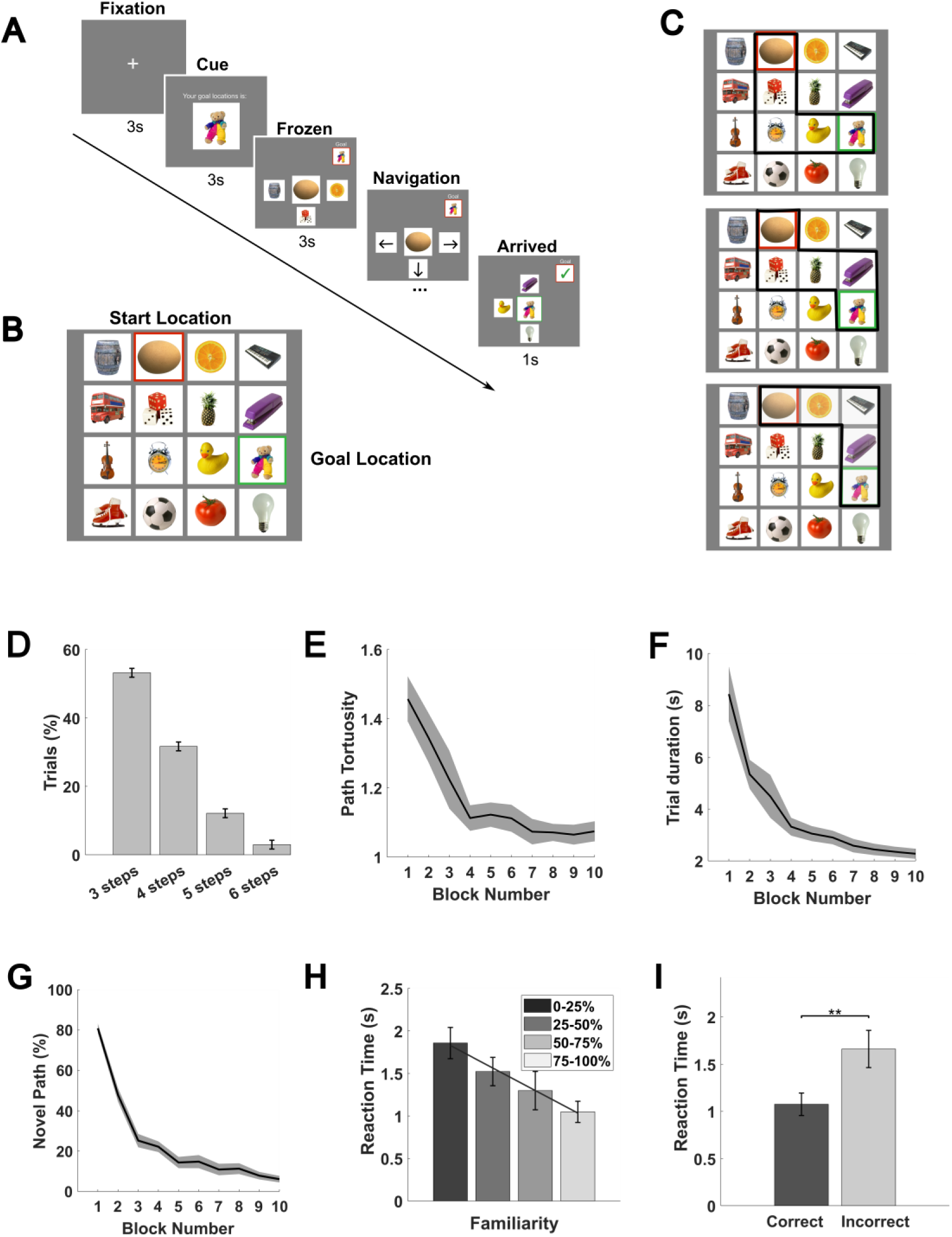
Schematic illustration of task structure and behavioural data. **(A)** During each trial, participants were cued with their goal location for 3s (‘cue period’) after a 3s fixation period and then frozen at their starting location on the abstract image map for 3s (‘frozen period’) before being allowed to navigate freely. During navigation, participants could use four buttons to move directly up, down, left or right from their current location on the abstract image map. Participants were instructed to find the shortest path from their current location to the given goal location in each trial. **(B)** Example abstract image map composed of sixteen visual stimuli. **(C)** A subset of the potential shortest paths between start and goal locations shown in (B). There are often multiple shortest paths available between a given start and goal location. **(D)** Distribution of shortest possible path distance from start to goal across 100 trials per participant. **(E)** Changes in path tortuosity across blocks of ten trials. **(F)** Changes in time taken to complete each path by block. **(G)** Changes in the proportion of completely novel paths traversed by block. **(H)** Reaction time taken to make the first transition, binned by the proportion of transitions that were previously experienced on the subsequent path. **(I)** Reaction times for correct and incorrect trials. All plots show mean values and error bars indicate SEM across participants unless otherwise stated. **= *p*<0.01.

In each trial, participants were presented with a goal image for 3s (‘cue period’) after a 3s fixation period, and then placed on the map but frozen in their current location for another 3s (‘frozen period’) before being allowed to navigate freely (Fig. 1A). The goal location in each trial was chosen at random from all locations that were at least three steps away from the current location on the map, and the start location in each trial was always the goal location from the previous trial. In all trials apart from the first trial in each task block, therefore, participants knew the start and goal location they had to navigate between during the cue period. The goal image was also displayed in the top right-hand corner of the screen throughout navigation in each trial, changing to a green tick when the goal location was reached. Upon reaching the goal, the screen was frozen for 1s on the goal image before the start of the next trial. Each participant completed 100 trials in the MEG scanner across three task blocks.

### Behavioural Analysis

To evaluate participants’ performance, correct trials were defined as those in which the shortest possible path to the goal image was taken (noting that there are often multiple different shortest paths, Fig. 1C). Trial-by-trial performance was also characterised by path tortuosity, defined as the total number of steps taken divided by the length of the shortest possible path (equal to one for correct trials, and greater than one for incorrect trials). We define ‘novel’ paths as those in which at least one transition had not been experienced before in either direction. Thereafter, the familiarity of each path is quantified as the number of transitions on that path that have been previously experienced divided by the total number of transitions made. Reaction times are defined as the time taken to make the first button press after the end of the frozen period.

### MEG Acquisition and Pre-processing

MEG data were recorded at a sample rate of 600 Hz using a whole-head 275-channel axial gradiometer system (CTF-Omega, VS Med Tech), while participants sat upright in a magnetically shielded room. Head position coils were attached to the nasion, left and right preauricular sites for anatomical co-registration.

MEG data were pre-processed using SPM12 (Litvak et al., 2011), Fieldtrip (Oostenveld et al., 2011), and MATLAB custom code. First, data were imported into MATLAB and cropped to the start and end of the task period. Data were then high-pass (0.1Hz) and notch (48-52Hz) filtered to remove slow drift and mains noise, respectively. Physiological artefacts related to eye blinks and lateral eye movements were identified and removed using independent components analysis implemented in Fieldtrip and EEGLAB (Delorme and Makeig, 2004). Two bad channels were identified in recordings from all participants using a generalised extreme studentised deviate test adopted from the OHBA software library (<u>OSL</u>) and removed from all subsequent analyses. Finally, data from each trial were ‘epoched’ into shorter time windows: from [−5 8s] around the onset of each cue period (‘spatial planning’ epochs); and [−3 3s] around the onset of each image during subsequent movement across the map (‘navigation’ epochs). Data from each participant were then merged across task blocks.

### Time-frequency Analysis

Estimates of dynamic changes in oscillatory power during task periods of interest were generated using SPM12. Specifically, power estimates were obtained for 20 logarithmically spaced frequency bands in the 1-30Hz range by convolving the signal with a five-cycle Morlet wavelet. Power values were log-transformed and then baseline-corrected before being submitted to subsequent analyses. Time-frequency data for the 13s ‘spatial planning’ epochs were baseline corrected using mean power in each frequency band during a [−1.5 −0.5s] window prior to the onset of the cue period (i.e. corresponding to a [1.5 2.5s] windows during the preceding fixation period) and then cropped to the [0 3s] cue period to avoid edge effects. Time-frequency data for the 6s ‘navigation’ epochs were baseline corrected using mean power in each frequency band during a [−1 −0.5s] window prior to each button press and then cropped to a [−1 1s] window around each button press to avoid edge effects. Navigation epochs in which two buttons were pressed simultaneously were excluded from subsequent analysis.

### MEG Source Reconstruction

MEG source reconstruction was conducted using the linearly constrained minimum variance (LCMV) beamformer (Barnes and Hillebrand, 2003) in SPM12 with a single-shell forward model to generate maps of mean source power on a 10-mm grid co-registered to MNI coordinates. As the LCMV beamformer requires a baseline period with a duration equal to the time window of interest, the 3s fixation period prior to cue onset was used as a baseline for cue period power contrasts, while no baseline was selected for the navigation period. In some cases, results were small volume corrected using anatomical masks generated by the WFU PickAtalas (Maldjian et al., 2003) with hippocampal regions defined by the Automated Anatomical Labelling Atlas (Tzourio-Mazoyer et al., 2002), or probabilistic masks of the entorhinal cortex from previous studies thresholded at 40% (Amunts et al., 2020, Convertino et al., 2023).

### MEG Source Regression

To identify relationships between behaviour and oscillatory power in source space, we conducted linear regression between the trial-by-trial power estimates in each voxel obtained by source localisation and distance to the goal using the multiple linear regression module implemented in SPM. Importantly, power estimates for both the cue and navigation periods were not baseline-corrected in this case. In addition, we included the trial number as a nuisance regressor to remove any potential effect of slow signal drift. Images of regression coefficients across voxels were then submitted to a one-sample t-test across participants with an uncorrected threshold of *p*<0.001, and results were reported if they passed a family-wise error (FWE) correction threshold of *p*<0.05 at the cluster level, either across the whole brain or following small volume correction.

### Phase-amplitude Coupling Analysis

To examine phase-amplitude coupling (PAC) between 2-5 Hz theta phase and the amplitude of slow and fast gamma bands (30-70 Hz and 70-140 Hz, respectively), we first reconstructed the filtered signal in each frequency band of interest in each source space voxel using weights from the LCMV beamformer and a zero phase, finite impulse response filter. Next, we extracted the analytical signal using the Hilbert transform and computed the resultant vector length of theta phase weighted by instantaneous gamma amplitude across all time bins in each voxel for each window of interest (Canolty et al., 2006, Tort et al., 2010). To remove any potential confound caused by concurrent trial-by-trial changes in oscillatory power, we constructed a general linear model to predict phase-amplitude coupling from theta and gamma power in each trial and then generated trial-by-trial images of the residuals to enter into our regression analysis against distance to the goal with trial number as a nuisance regressor. Finally, images of regression coefficients across voxels were submitted to a one-sample t-test across participants with an uncorrected threshold of *p*<0.001, and results were reported if they passed a family-wise error (FWE) correction threshold of *p*<0.05 at the cluster level, either across the whole brain or following small volume correction.

## Results

### Behavioural data

We asked 23 participants to complete an abstract navigation task inside the MEG scanner. In this task, participants were asked to move across a fixed 4 x 4 grid of images to find a specific goal image in each trial using as few steps as possible (Fig.1A-D). Participants rapidly learned the map layout and could subsequently navigate efficiently from start to goal locations in each trial. In particular, both path tortuosity (i.e. the total number of steps taken divided by the shortest possible path, Fig. 1E) and trial duration (time taken to reach the goal during navigation, Fig. 1F) decreased rapidly over the course of the task as participants came to experience every possible transition on the map (Fig.1G). The time taken to begin moving across the map also decreased with familiarity of the upcoming path (Fig.1H), suggesting the participants took less time to plan familiar routes (t(22) = −6.04, *p* < 0.001, one-sample t-test). Similarly, the time taken to execute each button press during navigation decreased with prior experience of each transition (t(22) = −7.06, *p* < 0.001, one-sample t-test). In total, participants took the ‘correct’ (i.e. shortest) path in 86.7 ± 2.18% of trials, and reaction times in those trials were slightly but significantly faster than in incorrect trials (t(22) = −3.71, *p* = 0.0012, paired t-test; Fig 1I).

### Hippocampal theta power during planning encodes goal distance

Previous intracranial EEG studies have suggested that theta power in the human hippocampal formation increases with distance to a hidden goal during navigation (Bush et al., 2017, Liu et al., 2023). Similarly, frontal theta power has been shown to increase with working memory load (Jensen and Tesche, 2002). To establish whether we could observe a similar oscillatory signature during spatial planning, we examined the spectral content of the MEG signal collapsed across all sensors during the 3s cue period (when the goal location for that trial was displayed on screen). We observed an increase in 2-5Hz theta power relative to a preceding baseline period that peaked during the first 1s (Fig. 2A). As expected, the resultant power spectrum (averaged over the full 3s cue period) also showed an increase in theta power versus baseline (Fig. 2B). To identify the source of this signal, we source localized 2-5Hz theta power during the full 3s window and found a large cluster centred on the prefrontal cortex ([−4, 24, −8]; Z = 6.60, *p*_FWE_ = 0.005, Fig. 2C) that extended into the bilateral medial temporal lobes ([6, −18, −44]; Z=4.34, *p*_FWE_ = 0.022; Supplementary Fig. 1A) and hippocampi ([−32, −20, −20], Z = 4.74, *p_FWE, SVC_* < 0.05; Supplementary Fig. 1B).

**Figure 2:**
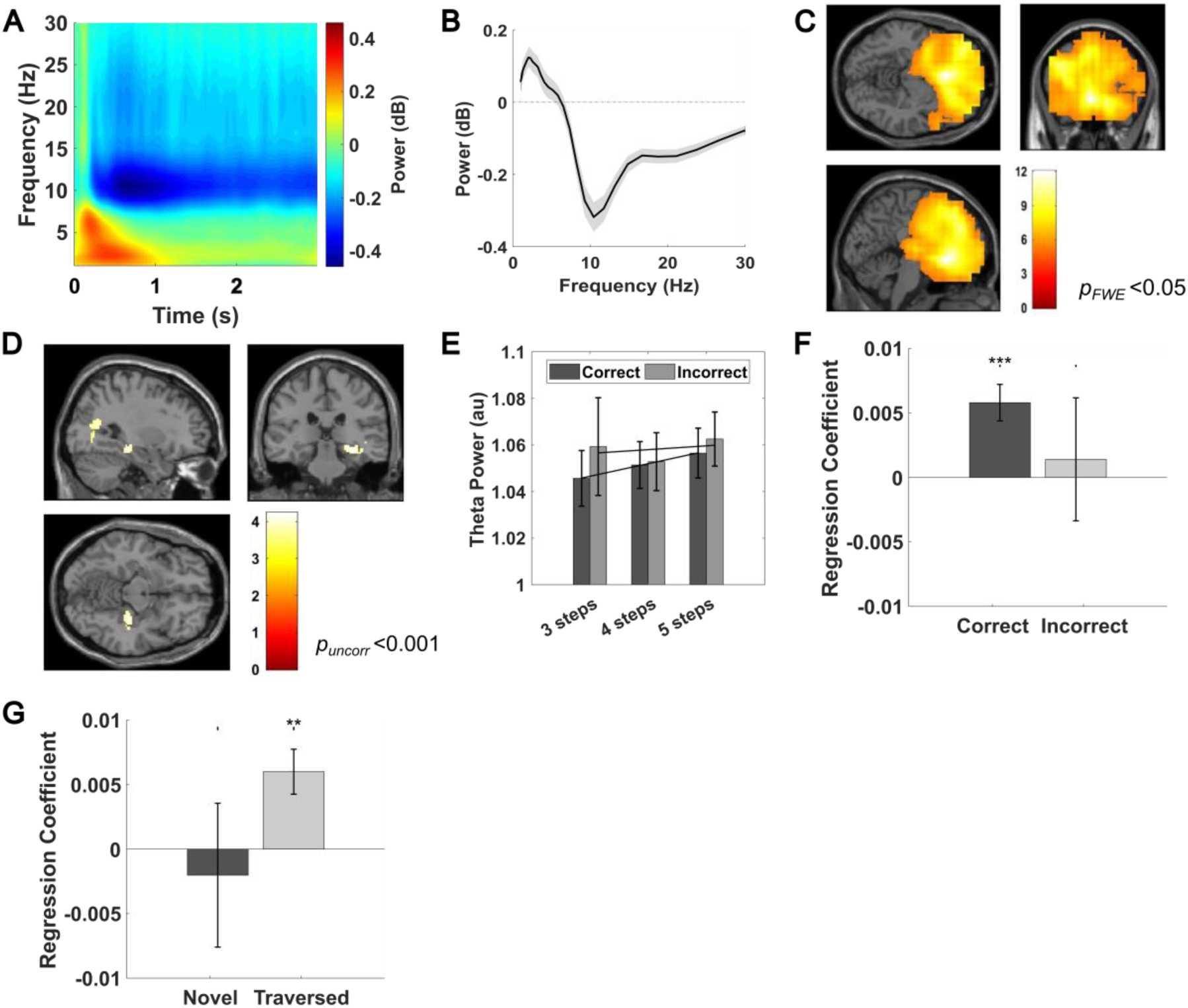
Theta power covaries with distance to the goal during planning. **(A)** Spectrogram of low-frequency oscillatory power during the 3s cue period, baseline corrected by mean power during 1s of the preceding fixation period. **(B)** Average power spectrum during the 3s cue period, baseline corrected by mean power during 1s of the preceding fixation period. **(C)** Source localisation of 2-5Hz theta power during the 3s cue period (visualised at *p_FWE_* < 0.05). Four clusters showed a significant increase in theta power ([−4, 24, −8]; Z = 6.6, *p_FWE_* < 0.001; [46, - 18, −44], Z = 4.34, *p_FWE_* = 0.005; [60, −30, 6]; Z = 4.13, *p_FWE_* = 0.046; [12, −2, 8], Z = 4.12, *p_FWE_* = 0.048). **(D)** Source localisation of 2-5Hz theta power during the 3s cue period that covaries positively with the shortest path distance to the goal (visualised at *p_uncorr_* <0.001). **(E)** Mean 2-5Hz theta activity extracted from a pre-defined right hippocampal ROI across the 3s cue period for trials with different shortest path lengths, shown separately for correct and incorrect trials. **(F)** Mean regression coefficient of 2-5Hz theta activity extracted from the right hippocampal ROI against distance to goal in correct (t(22) = 4.11, *p* <0.001, one-sample t-test) and incorrect (t(22) = 0.203, *p*= 0.843, one-sample t-test) trials. No significant difference is observed between correct and incorrect trials (t(22) = 0.713, *p* = 0.492, paired t-test). **(G)** Mean regression coefficient of 2-5Hz theta activity extracted from the right hippocampal mask against distance to goal for the novel (t(22) = −0.362, *p* = 0.722, one-sample t-test) and previously traversed (t(22) = 3.44, *p* = 0.0023, one-sample t-test) paths. No significant difference is observed between novel and traversed trials (t(22) = −1.52, *p* = 0.148, paired t-test). The colour bar in panels **C** and **D** shows t-statistics. All plots show mean values and error bars indicate SEM across participants unless otherwise stated. ***= *p*<0.001.

Next, to examine potential behavioural correlates of this oscillatory signal, we covaried theta power with step distance to the goal location during ‘correct’ trials (where participants knew the shortest path to the goal), separately for each source voxel. This revealed a significant cluster over the right medial temporal lobe ([30, −28, 10]; Z = 3.63, *p_uncorr_* <0.001, Fig. 2D) which passed our cluster-level statistical threshold within a pre-defined right hippocampal ROI ([30, −28, 10]; Z = 3.63, *p_FWE, SVC_* = 0.005; Supplementary Fig. 1C). No significant clusters were identified when the analysis was repeated using data from incorrect trials, either on a whole-brain level or within the same right hippocampal ROI. To further explore this effect, we extracted average theta power from the pre-defined right hippocampal ROI separately for correct and incorrect trials with 3, 4 and 5-step shortest paths to the goal (Fig. 2E). This illustrates a significant decrease in theta power with shortest path length from start to goal locations for correct trials (t(22) = 4.11, *p* < 0.001, one-sample t-test; Supplementary Fig. 1D), but no overall change for incorrect trials (where the participants did not know the shortest path to the goal, Fig. 2F).

Next, we aimed to establish whether the coding of goal distance in theta band oscillations arose equally from the planning of novel routes and the retrieval of previously executed paths across the map. To do so, we performed the same linear regression analysis between trial-by-trial theta power extracted from the right hippocampal ROI and distance to the goal, splitting correct trials into novel paths (where at least one transition had not been experienced before, in either direction) and previously traversed paths (where all transitions had been experienced before). Interestingly, a significant relationship between theta activity and goal distance was only observed for previously traversed paths (t(22) = 3.44, p = 0.0023, one-sample t-test; Fig. 2G). This suggests that the theta power code for goal distance during spatial planning might primarily reflect the retrieval of previously experienced navigational trajectories.

### Hippocampal theta power during navigation encodes goal distance

Next, we asked whether the relationship between theta power and distance to the goal would persist throughout navigation, as suggested by previous intracranial EEG studies (Liu et al., 2023). To do so, we first examined changes in oscillatory power collapsed across all MEG sensors on a movement-by-movement basis as participants navigated across the map using a button box. We observed an increase in 2-9 Hz theta power in a 2s window centred on each button press compared with a baseline period when participants were stationary on the map that peaked in a ∼1s window around translational movement (Fig. 3A). As expected, the resultant power spectrum (averaged over the full 2s period) also showed an increase in theta power versus baseline (Fig. 3B). Next, we extracted average power in low (2-5 Hz) and high (6-9 Hz) theta bands across all sensors during the 2s period around each button press during navigation (excluding the final step, when the goal is visible on screen). For correct trials (where participants knew where they were going), average power across all sensors in both low (t(22) = 6.28, p< 0.001, one-sample t-test) and high (t(22) = 6.36, *p* <0.001, one-sample t-test) theta bands iteratively decreased as the goal was approached, but this effect was less consistent for incorrect trials, where participants failed to take their shortest route to the goal (Fig. 3C; Supplementary Fig. 2).

**Figure 3:**
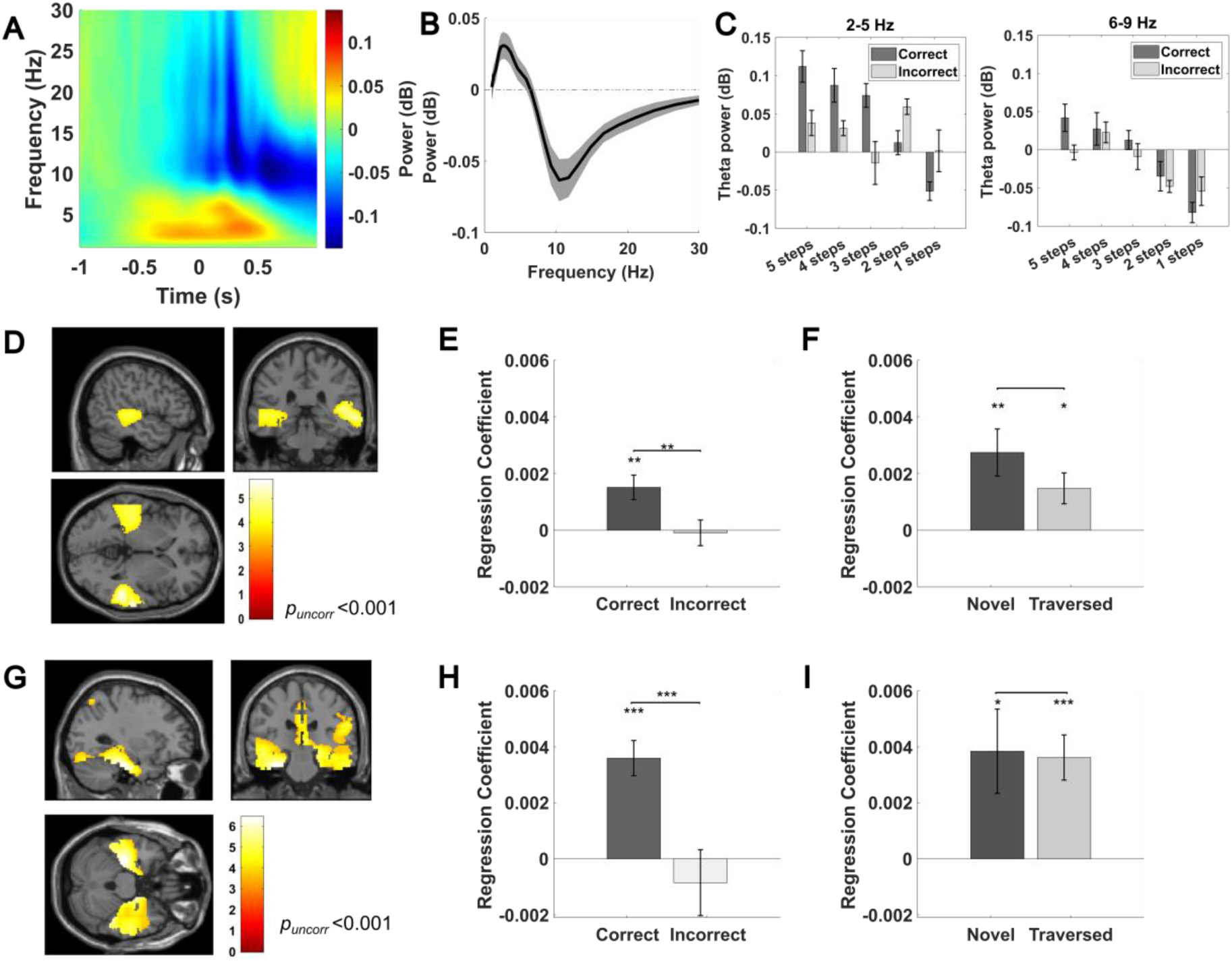
Theta power covaries with distance to the goal during navigation. **(A)** Spectrogram of low-frequency oscillatory power during 2s epochs centred on each button press during navigation, baseline corrected by mean power during the first 0.5s of the 2s epoch around each button press. **(B)** Average power spectrum during 2s button press epochs, baseline-corrected by mean power during the first 0.5s of that epoch. **(C)** Average 2-5 Hz and 6-9 Hz theta power across all sensors for correct and incorrect trials, split by distance to the goal in descending order during navigation. **(D)** Source localisation of the 2-5Hz theta power vs goal distance relationship (visualised at *p_uncorr_* < 0.001 without masking). Two clusters passed our cluster level significance threshold within a bilateral hippocampal mask (left: [−34, −34, −2]; Z = 3.43; *p_FWE, SVC_* = 0.026; right: [36, −14, −24]; Z = 3.14, *p_FWE, SVC_* = 0.042) **(E)** Mean regression coefficients for average 2-5Hz theta power extracted from a bilateral hippocampal mask against distance to the goal for correct and incorrect trials. **(F)** Mean regression coefficients for average 2-5Hz theta power extracted from a bilateral hippocampal mask against distance to the goal for novel and traversed trials. No significant difference is observed (t(22) = 1.34, *p* = 0.195, paired t-test). (G) Source localisation of the 6-9Hz theta power vs goal distance relationship (visualised at *p_uncorr_* < 0.001 without masking). Two clusters passed our cluster-level significance threshold within a bilateral hippocampal mask (left: [− 34, −26, −16]; Z = 4.15; *p_FWE, SVC_* = 0.012; right: [16, −10, −18]; Z = 4.47; *p_FWE, SVC_* = 0.005). **(H)** Mean regression coefficients for average 6-9Hz theta activity extracted from a bilateral hippocampal mask against distance to the goal for correct and incorrect trials trials. **(I)** Mean regression coefficients for average 6-9Hz theta power extracted from a bilateral hippocampal mask against distance to the goal for novel and traversed paths. No significant difference is observed (t(22) = 0.127, *p* = 0.90, paired t-test). The colour bar in panels **D** and **H** shows t-statistics. All plots show mean values and error bars indicate SEM across participants unless otherwise stated. ** = *p* <0.01, *** = *p* <0.001.

To identify the source of this signal, we covaried step distance to the goal location during ‘correct’ trials (where the participants knew the shortest path to reach the goal) with theta power in each source voxel. This revealed two significant clusters centred over the bilateral temporal lobes for both low ([52, −38, 4]; Z = 4.45, *p_FWE_* = 0.017, Fig. 3D) and high ([−30, −24, −36]; Z = 4.77, *p_FWE_* = 0.005; Fig. 3G) theta bands, each of which passed our cluster level statistical threshold within a pre-defined bilateral hippocampal ROI (low theta: [−34, −34, −2]; Z = 3.43, *p_FWE, SVC_* < 0.05; Supplementary Fig. 3A; high theta: [16, −10, −18]; Z = 4.77, *p_FWE, SVC_* < 0.05; Supplementary Fig. 4A). To further explore this effect, we extracted average theta power from the same bilateral hippocampal ROI separately for correct and incorrect trials with different numbers of steps remaining to the goal location. For correct trials, we found a significant positive relationship between theta power in the bilateral hippocampus and distance remaining to the goal in both the 2-5 Hz (t(22) = 3.51, *p*= 0.002, one-sample t-test; Fig. 3E and Supplementary 3B) and 6-9 Hz (t(22) = 5.71, *p*< 0.001, one-sample t-test; Fig. 3H and Supplementary 4B) frequency bands. In both cases, theta power decreased iteratively as the goal was approached in correct trials only (Supplementary Fig. 3C-E; 4C), and the relationship between theta power and distance to the goal was stronger in correct versus incorrect trials (low theta: t(22) = 3.37, *p* = 0.0028, paired t-test; Fig. 3E; high theta: t(22) = 4.10, *p* < 0.001, paired t-test; Fig. 3H).

Finally, we sought to confirm whether the theta power code for goal distance arose primarily during the navigation of previously traversed routes, as observed during the cue period. During navigation, however, we observed a significant positive relationship between bilateral hippocampal 2-5Hz theta power and distance remaining to the goal for both novel (t(22) = 3.31, p = 0.0032, one-sample t-test) and previously traversed paths (t(22) = 2.72, *p* = 0.0124, one-sample t-test; Fig. 3F). Similarly, a significant positive relationship between bilateral hippocampal 6-9 Hz theta power and distance to the goal was present for both novel (t(22) = 2.55, *p* = 0.0181, one-sample t-test) and previously traversed paths (t(22) = 4.49, *p* < 0.001, one-sample t-test; Fig. 3I). This suggests that hippocampal theta power distance to goal coding during active navigation does not depend on prior experience of the route being traversed.

### Theta-gamma phase-amplitude coupling mediates spatial planning

It has been hypothesised that the phase of theta oscillations may modulate the amplitude of concurrent gamma band activity to organise sequential representations during memory function (Lisman and Jensen, 2013). According to this model, the encoding of longer sequences corresponds to a wider theta phase distribution of gamma power, which should therefore be negatively correlated with the strength of theta-gamma phase-amplitude coupling (TG-PAC; (Heusser et al., 2016, Daume et al., 2024). Hence, we examined whether TG-PAC would increase as the goal was approached and an increasingly short sequence of upcoming locations needed to be maintained in memory.

First, we examined dynamic changes in TG-PAC between 2-5 Hz theta phase and 70-140 Hz fast gamma amplitude as the goal is approached during navigation. To do so, we covaried step distance to the goal with TG-PAC in each source space voxel (excluding the final step, when the goal is visible on screen). This revealed a significant cluster in the right anterior temporal lobe for correct trials ([16, - 10,-30]; Z = 3.92, *p_uncorr_* < 0.001, Fig. 4A), which passed our cluster level statistical threshold in a pre-defined right entorhinal cortex ROI ([18, −10, −30]; Z = 3.91, *p_FWE, SVC_* = 0.004; Supplementary Fig. 5A). To further explore this effect, we extracted estimates of TG-PAC from the right entorhinal ROI and conducted a linear regression against distance remaining to the goal. We observed the same negative correlation between TG-PAC and distance to the goal in correct (t(22) = −2.90, *p* = 0.0082, one-sample t-test) but not incorrect (t(22) = −0.34, *p* = 0.74, one-sample t-test) trials (Fig. 4B and 4C; Supplementary Fig 5B). Intriguingly, we found that this relationship between theta–fast gamma phase-amplitude coupling and distance to the goal during navigation was present for novel (t(22) = −3.32, *p* = 0.0031, one-sample t-test; Fig. 4D, E and F) but not previously traversed paths (t(22) = −1.88, *p* = 0.0733, one-sample t-test). Consistent with this, covarying step distance to the goal during novel trials with TG-PAC in each source space voxel revealed a significant cluster in right entorhinal cortex ([18, - 6, −30]; Z = 3.77, *p_uncorr_* < 0.001) which passed our cluster-level statistical threshold in a pre-defined right entorhinal cortex ROI ([22, −18, −30]; Z = 3.77, *p_FWE, SVC_* = 0.006; Supplementary Fig. 5C). This is consistent with previous rodent electrophysiology studies that describe increased phase-amplitude coupling between theta and fast gamma band activity originating in the entorhinal cortex during the encoding of new information (Colgin et al., 2009).

**Figure 4:**
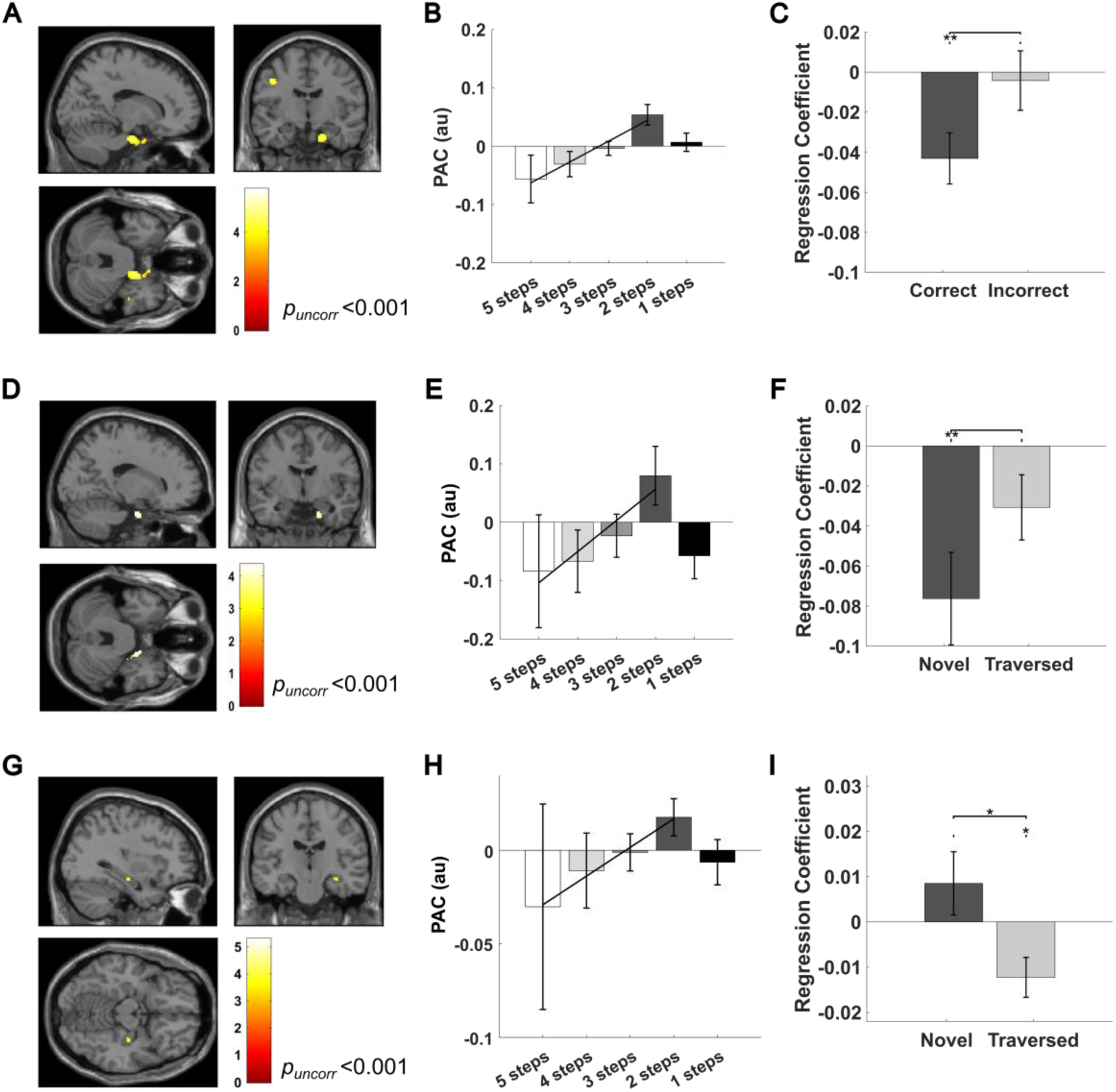
Theta-gamma phase-amplitude coupling covaries with distance to the goal during navigation. **(A)** Source localisation of the 2-5Hz theta - 70-140 Hz fast gamma phase-amplitude coupling vs goal distance relationship for all correct trials (visualised at *p_uncorr_* < 0.001). A significant cluster is observed in the right anterior temporal lobe. **(B)** Average theta-fast gamma PAC extracted from a pre-defined right entorhinal mask for all correct trials, split by distance to the goal in descending order during navigation. **(C)** Regression coefficients between theta-fast gamma PAC and goal distance in the right entorhinal mask for correct and incorrect trials (excluding the final step, when the goal is visible on screen). No significant difference is observed (t(22) = −1.65, *p* = 0.11, paired t-test). **(D)** Source localisation of the theta–fast gamma PAC vs goal distance relationship for all correct trials that use novel paths (visualised at *p_uncorr_* < 0.001). **(E)** Average theta-fast gamma PAC extracted from a pre-defined right entorhinal mask for all correct trials that use novel paths, split by distance to the goal in descending order during navigation. **(F)** Regression coefficients between theta-fast gamma PAC and goal distance in the right entorhinal mask, for all correct trials that use novel and previously traversed paths (excluding the final step, when the goal is visible on screen). No significant difference is observed (t(22) = −1.77, *p* = 0.119, paired t-test). **(G)** Source localisation of the 2-5Hz theta - 30-70 Hz slow gamma phase-amplitude coupling vs goal distance relationship for all correct trials that use previously traversed paths (visualised at *p_uncorr_* < 0.001). (H) Average theta-slow gamma PAC extracted from a pre-defined right hippocampal mask for all correct trials that use previously traversed paths, split by distance to the goal in descending order during navigation. **(I)** Regression coefficients between theta-slow gamma PAC and goal distance in the right hippocampal mask for all correct trial that use novel and previously traversed paths (excluding the final step, when the goal is visible on screen). The colour bar in panels **A, D,** and **G** shows t-statistics. All plots show mean values and error bars indicate SEM across participants unless otherwise stated. * = *p* <0.05, ** = *p* <0.01, *** = *p* <0.001.

Next, we examined TG-PAC between 2-5 Hz theta phase and 30-70 Hz slow gamma amplitude. Although we did not observe any significant clusters at the whole brain level (Supplementary Fig. 6A), we identified a cluster in the right hippocampus that showed a negative relationship between TG-PAC and distance to the goal during the traversal of previously experienced paths ([34, −18, −14]; Z = 3.25, *p_uncorr_* = 0.001; Fig. 4G), which is marginally significant in a pre-defined right hippocampal ROI ([34, - 18, −14]; Z = 3.25, *p_FWE, SVC_* = 0.051; Supplementary Fig. 6B). To further explore this effect, we conducted the same regression analysis between trial-by-trial estimates of TG-PAC obtained from the right hippocampal ROI and distance to the goal (Fig. 4H). This showed a significant negative relationship for previously traversed paths (t(22) = −2.78, p = 0.0109, one-sample t-test) but not for novel paths (t(22) = 1.25, *p* = 0.22, one-sample t-test), with the former being significantly stronger than the latter (t(22) = −2.09, *p* = 0.048, paired t-test, Fig. 4I; Supplementary Fig. 6C). This is consistent with previous rodent electrophysiology studies that describe increased phase-amplitude coupling between theta and slow gamma band activity originating within the hippocampus during the sequential reactivation of previously learned information (Colgin et al., 2009).

In summary, TG-PAC in the hippocampal formation increases dynamically as a hidden goal is approached during navigation. This is consistent with the hypothesis that maintaining a longer sequence of items in memory – i.e. a greater number of locations that remain to be traversed – corresponds to a wider distribution of gamma power in each theta cycle and, therefore, lower TG-PAC (Heusser et al., 2016, Daume et al., 2024). In addition, we found that distinct fast and slow gamma bands originating in the entorhinal cortex and within the hippocampus support the encoding and retrieval of new and previously traversed paths, respectively.

## Discussion

We have shown that human hippocampal theta power during both planning (immediately prior to movement onset) and active navigation correlates with the distance to a hidden goal location, only in correct trials when participants are aware of the distance they need to travel. This replicates previous intracranial electrophysiology studies in epilepsy patients (Bush et al., 2017, Liu et al., 2023) and extends those findings to a healthy population navigating in an abstract state space. Moreover, these results are analogous to fMRI studies showing that the hippocampal BOLD signal during navigation in naturalistic virtual environments correlates with path distance to a hidden goal, while the entorhinal BOLD signal correlates with Euclidean distance to the same goal (Howard et al., 2014). During the planning period, we also observed widespread increases in theta power across frontal cortex that did not covary with subsequent path length. Alongside a role in cognitive control and working memory function (Cavanagh and Frank, 2014), frontal midline theta oscillations have been implicated in spatial and episodic memory retrieval processes (Kaplan et al., 2014, Adams et al., 2020, Herweg et al., 2020). Interestingly, the modulation of hippocampal theta power by subsequent path distance during this period was strongest prior to the traversal of previously experienced paths. Hence, our results suggest that the frontal theta rhythm coordinates the retrieval of past experience during spatial planning to provide an estimate of goal distance that is subsequently reflected in hippocampal theta power.

These findings are also reminiscent of working memory studies in which frontotemporal theta power increases with the number of items that must be actively rehearsed for several seconds before retrieval (Jensen and Tesche, 2002, Tesche and Karhu, 2000, Sternberg, 1966). In our study, participants may be actively rehearsing the sequence of locations they intend to traverse to reach the goal. This would explain the dynamic reduction in hippocampal theta power as the goal is approached during navigation, and the number of locations remaining to be traversed decreases. Elsewhere, it has been hypothesised that sequences of memory representations are encoded by gamma band activity at successive theta phases during learning and maintenance periods(Jensen and Lisman, 1998, Lisman and Idiart, 1995, Lega et al., 2016). According to this view, theta-gamma phase amplitude coupling should decrease as sequence length increases, and gamma power becomes more widely distributed across the theta cycle (Heusser et al., 2016, Daume et al., 2024). Consistent with this, we observed a dynamic increase in theta-gamma phase-amplitude coupling as the goal was approached during navigation, and fewer locations remained to be traversed.

In addition, we found that theta phase modulated fast gamma amplitude in the entorhinal cortex during the encoding of new paths and slow gamma amplitude in the hippocampus during the retrieval of previously traversed paths. This replicates previous rodent (Colgin et al., 2009, Bieri et al., 2014, Zheng et al., 2016, Fernández-Ruiz et al., 2017, Dvorak et al., 2018, Lopes-dos-Santos et al., 2018) and human (Griffiths et al., 2019, Vivekananda et al., 2021) studies which have shown that fast and slow gamma power and phase-amplitude coupling originating from entorhinal cortex and intra-hippocampal projections are preferentially engaged during memory encoding and retrieval, respectively. Numerous rodent studies have also demonstrated how the hippocampal theta rhythm organises sequences of place cell firing in each oscillatory cycle to represent upcoming behavioural trajectories (O’Keefe and Recce, 1993, Burgess et al., 1994, Johnson and Redish, 2007, Skaggs et al., 1996), and that the length of spatial trajectories encoded by these theta sweeps is greater during navigation towards distant goals (Wikenheiser and Redish, 2015). As such, our findings are consistent with the hypothesis that theta sequences of upcoming locations in the human hippocampus are maintained by phase amplitude coupling, with fast gamma from the entorhinal cortex and slow gamma from intra-hippocampal sources dominating during memory encoding and retrieval, respectively.

In summary, we have demonstrated that hippocampal theta oscillations code for distance to a hidden goal during both spatial planning and subsequent navigation. In addition, we have shown that hippocampal theta modulation of gamma amplitude in distinct slow and fast bands increases iteratively as a hidden goal is approach during navigation of novel and previously traversed paths, respectively. This suggests that theta oscillations in the human hippocampal formation scaffold distance to goal coding during flexible navigation in an abstract state space.

## Acknowledgements

The authors wish to thank George O’Neill, Oliver Vikbladh, Danying Wang and all members of the UCL Human Electrophysiology Lab for their constructive feedback during the preparation of this manuscript, as well as the imaging support staff at the Wellcome Centre for Human Neuroimaging. This work was supported by a Wellcome Principal Research Fellowship to NB (222457/Z/21/Z) and a UKRI Frontier Research Grant to DB (EP/X023060/1).

---

**Figure S1.**
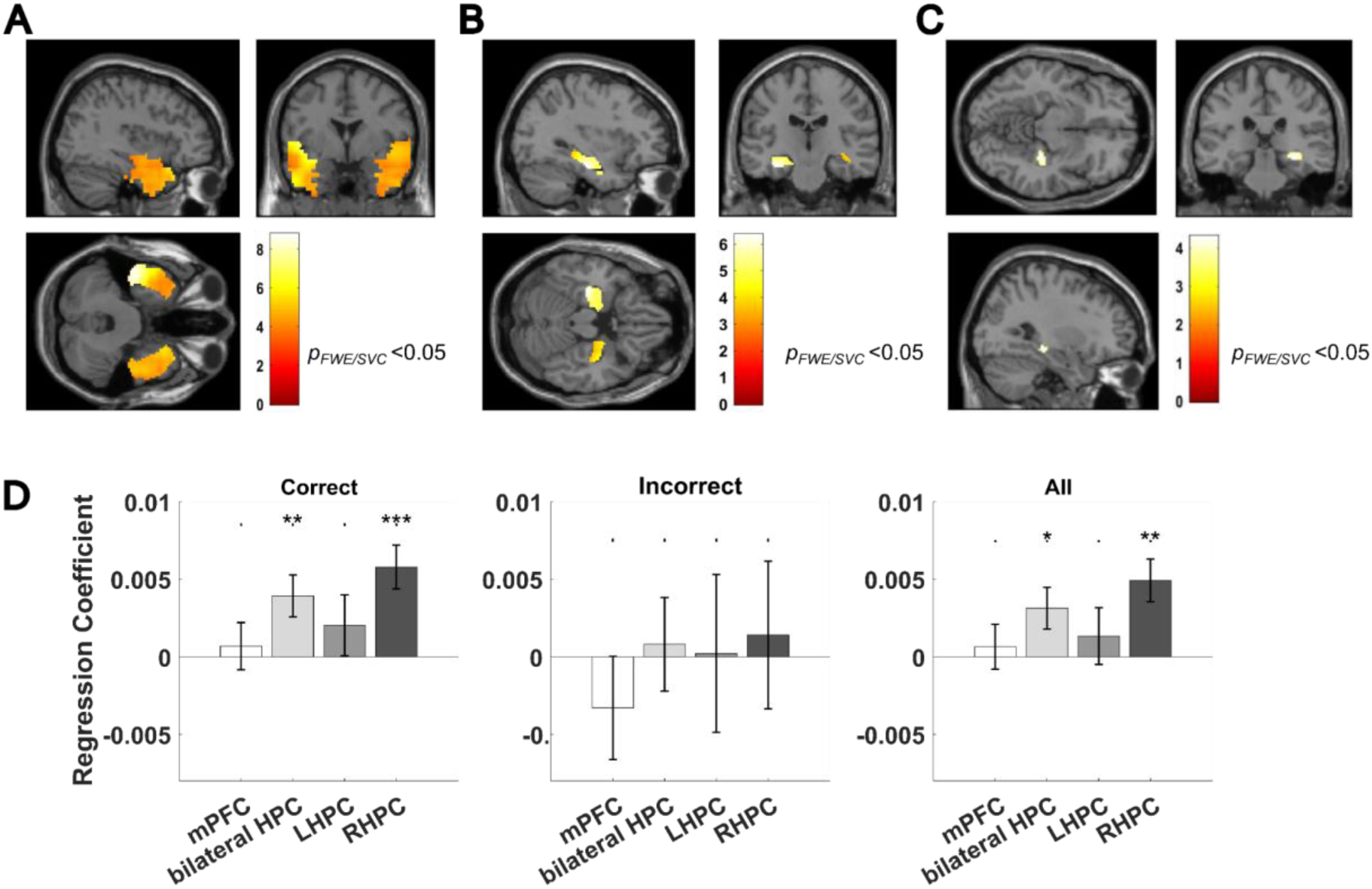
Frontotemporal theta power increases during spatial planning. **(A)** Source localisation of increase in 2-5 Hz theta power during the 3s cue period, masked and small volume corrected with a bilateral temporal lobe mask (visualised at *p_uncorr_* < 0.001). **(B)** Source localisation of increase in 2-5Hz theta power during the 3s cue period, masked and small volume corrected with a bilateral hippocampal mask (visualised at *p_uncorr_* <0.001). **(C)** Source localisation of the relationship between 2-5 Hz theta power during the 3s cue period and shortest path distance to the goal for all correct trials, masked and small volume corrected with a bilateral hippocampal mask (visualised at *p_uncorr_* <0.001). **(D)** Regression coefficients between 2-5Hz theta power during planning and shortest path distance to the goal extracted from pre-defined anatomical ROIs across correct trials (left panel), incorrect trials (middle panel), and all trials (right panel). Data shown for the medial prefrontal cortex (mPFC, correct trials: t(22) = 0.46, *p* = 0.653, one-sample t-test; incorrect trials: t(22) = −0.69, *p* = 0.508, one-sample t-test; all trials: t(22) = 0.453, *p* = 0.655, one-sample t-test), bilateral hippocampus (correct trials: t(22) = 2.89, *p* = 0.0085, one-sample t-test; incorrect trials: t(22) = 0.186, *p* = 0.856, one-sample t-test; all trials: t(22) = 2.33, *p* = 0.0291, one-sample t-test), right hippocampus (RHPC, correct trials: t(22) = 4.11, *p* < 0.001, one-sample t-test; incorrect trials: t(22) = 0.20, *p* = 0.843, one-sample t-test; all trials: t(22) = 3.57, *p* = 0.0017, one-sample t-test) and left hippocampus separately (LHPC, correct trials: t(22) = 1.03, *p* = 0.3118, one-sample t-test; incorrect trials t(22) = 0.029, *p* = 0.978, one-sample t-test; all trials: t(22) = 0.0727, *p* = 0.475, one-sample t-test). No significant difference is observed between correct trials and incorrect trials in any ROIs (all *p* > 0.36).

**Figure S2.**
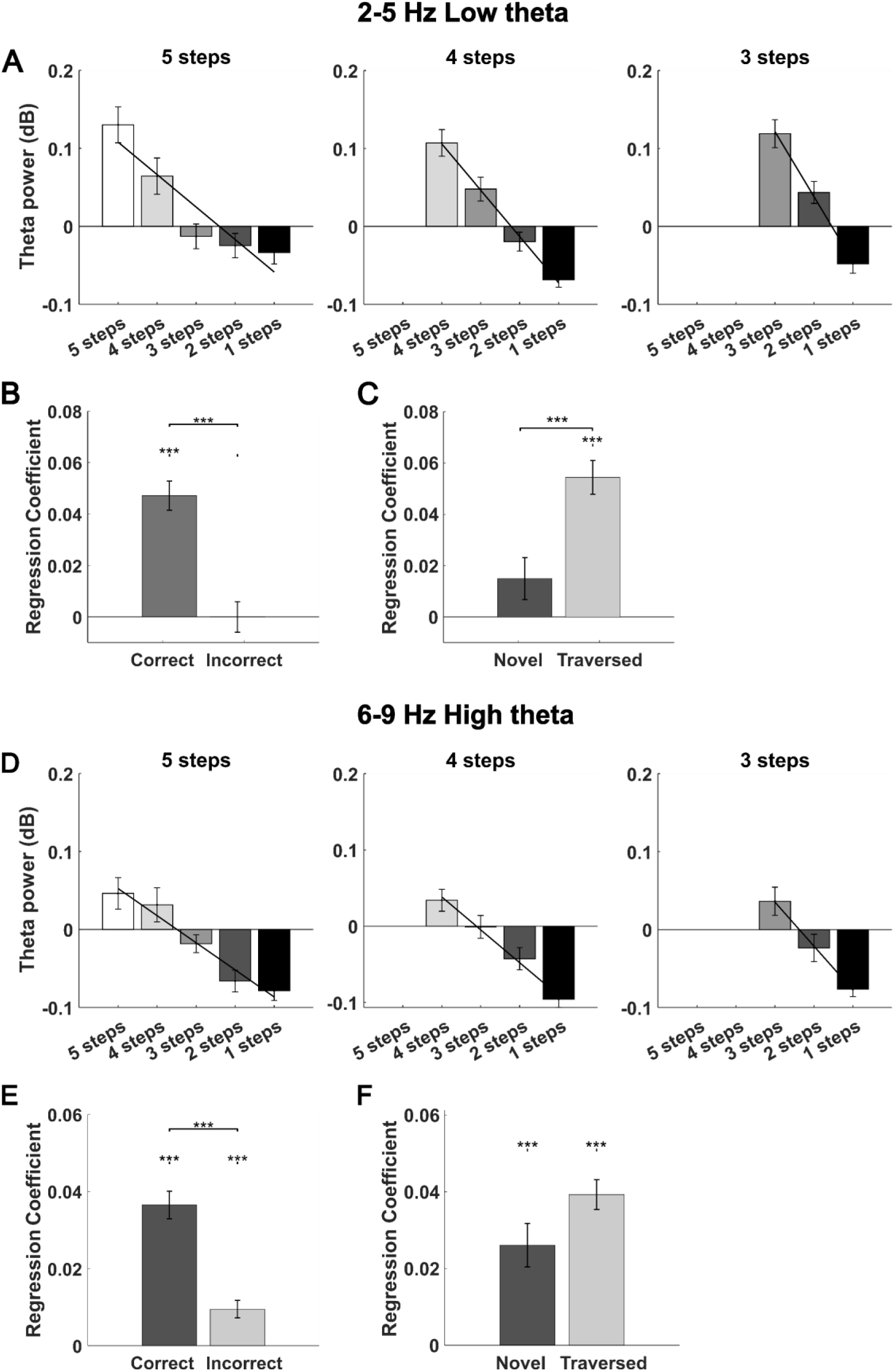
Scalp level theta power covaries with distance to the goal during navigation. **(A)** 2-5 Hz theta power averaged across all sensors for all correct trials, split by the remaining distance to the goal in descending order during navigation. Left panel: correct five-step paths; middle panel: correct four-step paths; right panel: correct three-step paths. **(B)** Regression coefficient for average 2-5 Hz theta power against distance to the goal for correct (t(22) = 6.28, p< 0.001, one-sample t-test) and incorrect trials (t(22) = −0.442, p = 0.662, one-sample t-test). This effect is significantly stronger in correct trials (t(22) = 4.97, *p* < 0.001, paired-t-test). **(C)** Regression coefficient for average 2-5Hz theta power against distance to the goal for correct trials that use novel (t(22) = 1.82, *p* = 0.0827, one-sample t-test) or previously traversed paths (t(22) = 8.33, *p* < 0.001, one-sample t-test). This effect is significantly stronger for previously traversed paths (t(22) = 4.39, *p* < 0.001, paired t-test). **(D)** 6-9 Hz theta power averaged across all sensors for all correct trials, split by the remaining distance to the goal in descending order during navigation. Left panel: correct five-step paths; middle panel: correct four-step paths; right panel: correct three-step paths. **(E)** Regression coefficient for average 6-9 Hz theta power against distance to the goal for correct (t(22) = 6.36, *p* < 0.001, one-sample t-test) and incorrect trials (t(22) = 3.02, *p* = 0.0062, one-sample t-test). This effect is significantly stronger in correct trials (t(22) = 4.23, *p* < 0.001, paired t-test). **(F)** Regression coefficient for average 6-9 Hz theta power against distance to the goal for correct trials that use novel (t(22) = 4.37, *p* < 0.001, one-sample t-test) or previously traversed paths (t(22) = 9.92, p < 0.001, one-sample t-test). The effect is marginally stronger for previously traversed paths (t(22) = 2.07, *p* = 0.0502, paired t-test).

**Figure S3.**
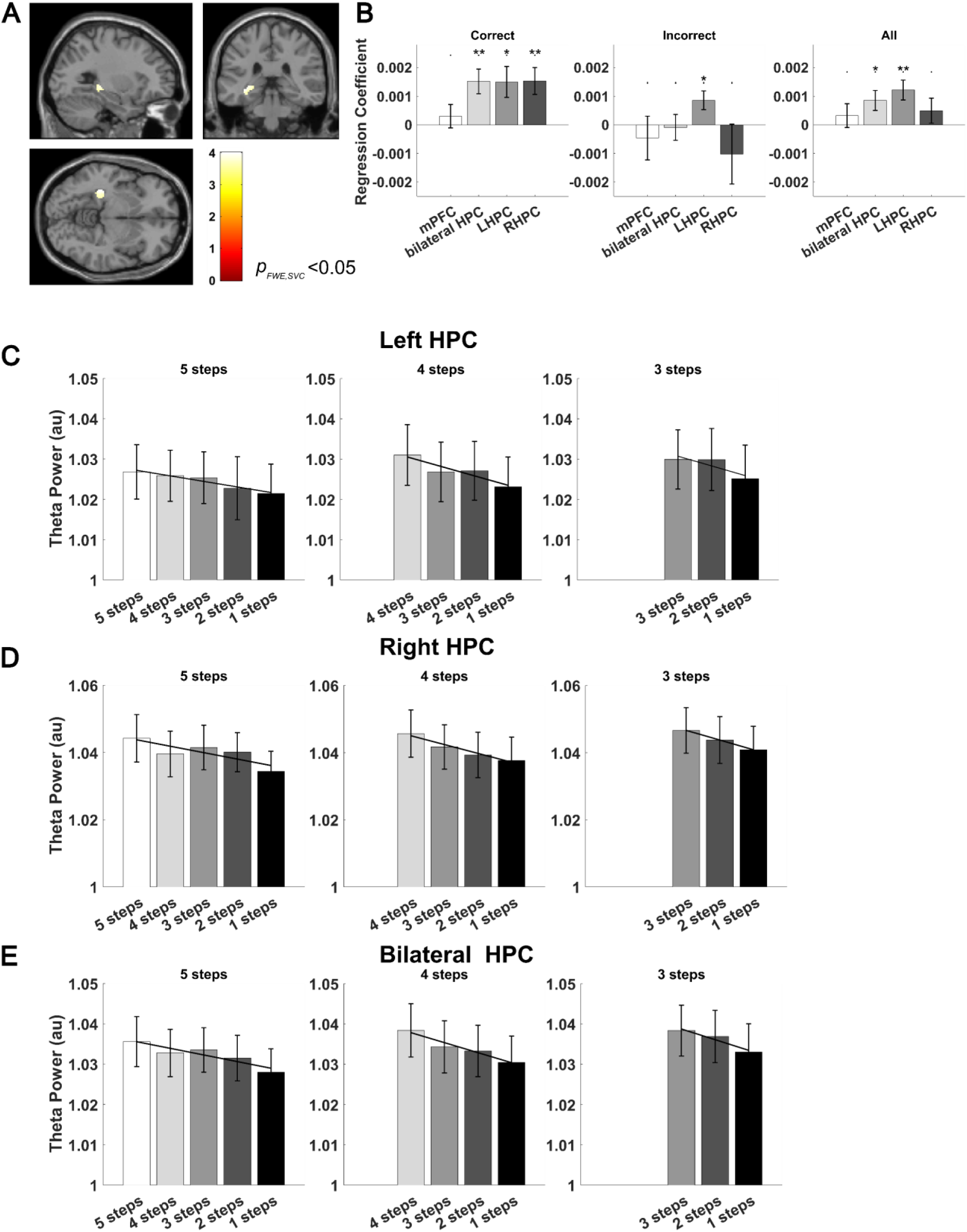
Source level 2-5Hz theta power covaries with distance to the goal during navigation. **(A)** Source localisation of the relationship between 2-5 Hz theta power and shortest path distance to the goal for all correct trials within a bilateral HPC mask (visualised at *p_uncorr_* < 0.001), with a peak in the left hippocampus ([−34, −34, −2]; Z = 3.43, *p_FWE, SVC_* < 0.05). **(B)** Regression coefficients between 2-5Hz theta power during navigation and shortest path distance to the goal extracted from pre-defined anatomical ROIs across correct trials (left panel), incorrect trials (middle panel), and all trials (right panel). Data shown for the medial prefrontal cortex (mPFC, correct trials: t(22) = 0.727, *p* = 0.475, one-sample t-test; incorrect trials: t(22) = −0.603, *p* = 0.551, one-sample t-test; all trials: t(22) = 0.77, *p* = 0.449, one-sample t-test), bilateral hippocampus (correct trials: t(22) = 3.51, *p* = 0.002, one-sample t-test; incorrect trials: t(22) = −0.20, *p* = 0.844, one-sample t-test; all trials: t(22) = 2.43, *p* = 0.0236, one-sample t-test), left hippocampus (LHPC, correct trials: t(22) = 2.76, *p* = 0.0114, one-sample t-test; incorrect trials: t(22) = 2.63, *p* = 0.0152, one-sample t-test; all trials: t(22) = 3.49, *p* = 0.0021, one-sample t-test) and right hippocampus (RHPC, correct trials: 3.26, *p* = 0.0036, one-sample t-test; incorrect trials: t(22) = −0.980, *p* = 0.338, one-sample t-test; all trials: t(22) = 1.12, *p* = 0.273, one-sample t-test). Distance to goal coding is significantly stronger in correct versus incorrect trials in the bilateral hippocampus (t(22) = 3.37, *p* = 0.0028, paired t-test) and right hippocampus (t(22) = 2.67, *p* = 0.014, paired t-test), while there is no significant difference in mPFC (t(22) = 1.08, *p* = 0.29, paired t-test) or left hippocampus (t(22) = 1.02, *p* = 0.317, paired t-test). **(C, D, E)** 2-5 Hz theta power extracted from (C) left hippocampus, (D) right hippocampus, and (E) bilateral hippocampus split by remaining distance to the goal in descending order during navigation. Left panel: correct five-step paths; middle panel: correct four-step paths; right panel: correct three-step paths.

**Figure S4:**
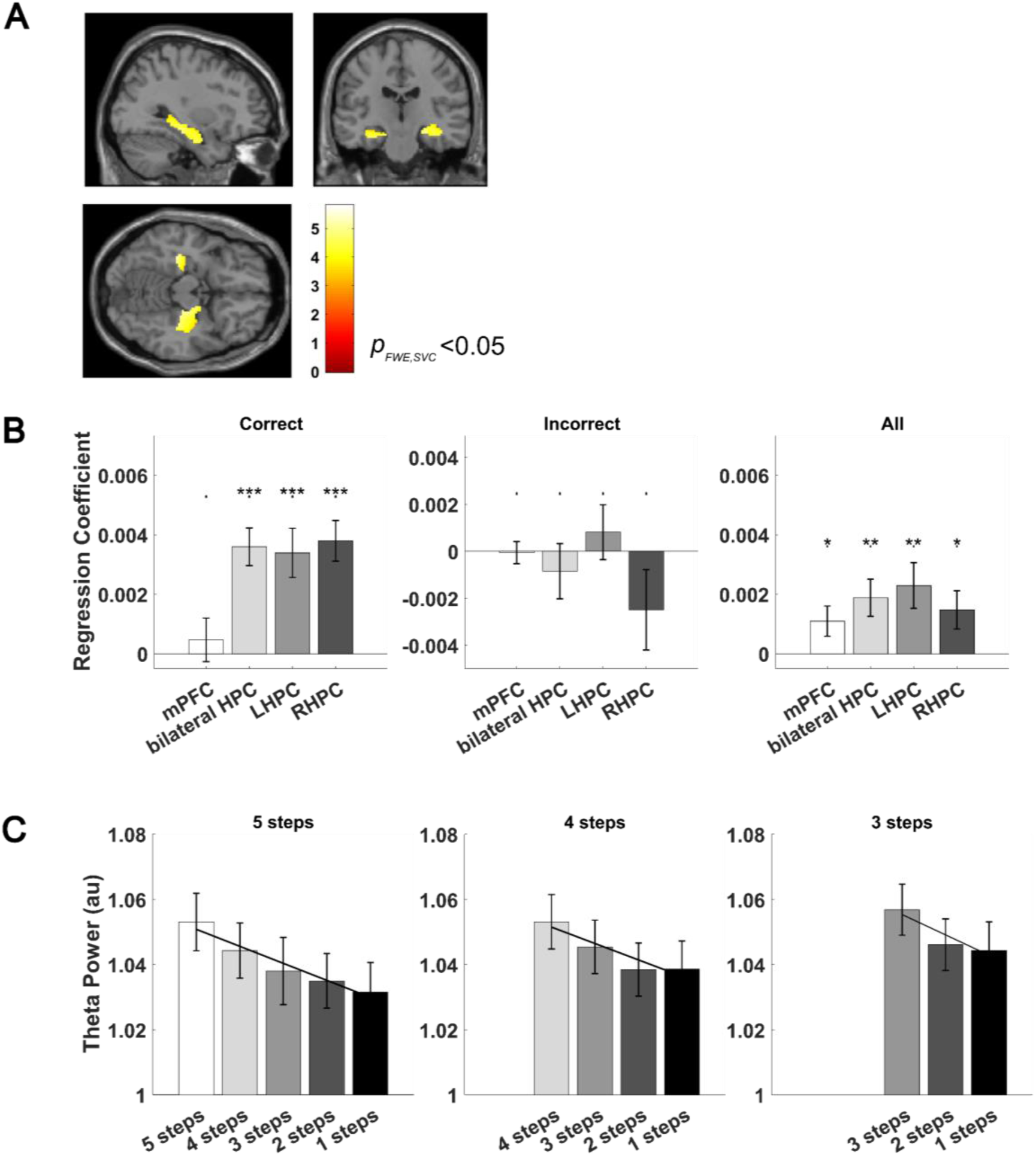
Source level 6-9Hz theta power covaries with distance to the goal during navigation. **(A)** Source localisation of the relationship between 6-9 Hz theta power and shortest path distance to the goal for all correct trials within a bilateral HPC mask (visualised at *p_uncorr_* < 0.001), with a global peak in the right hippocampus (16, - 10, −18; Z = 4.77, *p_FWE, SVC_* < 0.05). **(B)** Regression coefficients between 6-9Hz theta power during navigation and shortest path distance to the goal extracted from pre-defined anatomical ROIs across correct trials (left panel), incorrect trials (middle panel), and all trials (right panel). Data shown for the medial prefrontal cortex (mPFC, correct trials: t(22) = 0.636, *p* = 0.532, one-sample t-test; incorrect trials: t(22) = −0.128, *p* = 0.90, one-sample t-test; all trials: t(22) = 2.16, *p* = 0.0416, one-sample t-test), bilateral hippocampus (t(22) = 5.78, *p* < 0.001, one-sample t-test; incorrect trials: −0.724, *p* = 0.477, one-sample t-test; all trials: t(22) = 3.02, *p* = 0.0063, one-sample t-test), left hippocampus (LHPC, correct trials: t(22) = 4.10, *p* < 0.001, one-sample t-test; incorrect trials: t(22) = 0.693, *p* = 0.50, one-sample t-test; all trials: t(22) = 3.00, *p* = 0.0066, one-sample t-test) and right hippocampus (RHPC, correct trials: t(22) = 5.52, *p* < 0.001, one-sample t-test; incorrect trials: t(22) = −1.45, *p* = 0.161, one-sample t-test; all trials: t(22) = 2.28, *p* = 0.0325, one-sample t-test). Distance to goal coding is significantly stronger in correct versus incorrect trials in the bilateral hippocampus (t(22) = 4.10, *p* < 0.001, paired t-test), right hippocampus (t(22) = 3.53, *p* = 0.0029, paired t-test) and left hippocampus (t(22) = 2.60, *p* = 0.0164, paired t-test), while there is no significant difference in mPFC (t(22) = 0.578, *p* = 0.569, paired t-test). **(C)** 6-9 Hz theta power extracted from the bilateral hippocampus split by the remaining distance to the goal in descending order during navigation. Left panel: correct five-step paths; middle panel: correct four-step paths; right panel: correct three-step paths.

**Figure S5.**
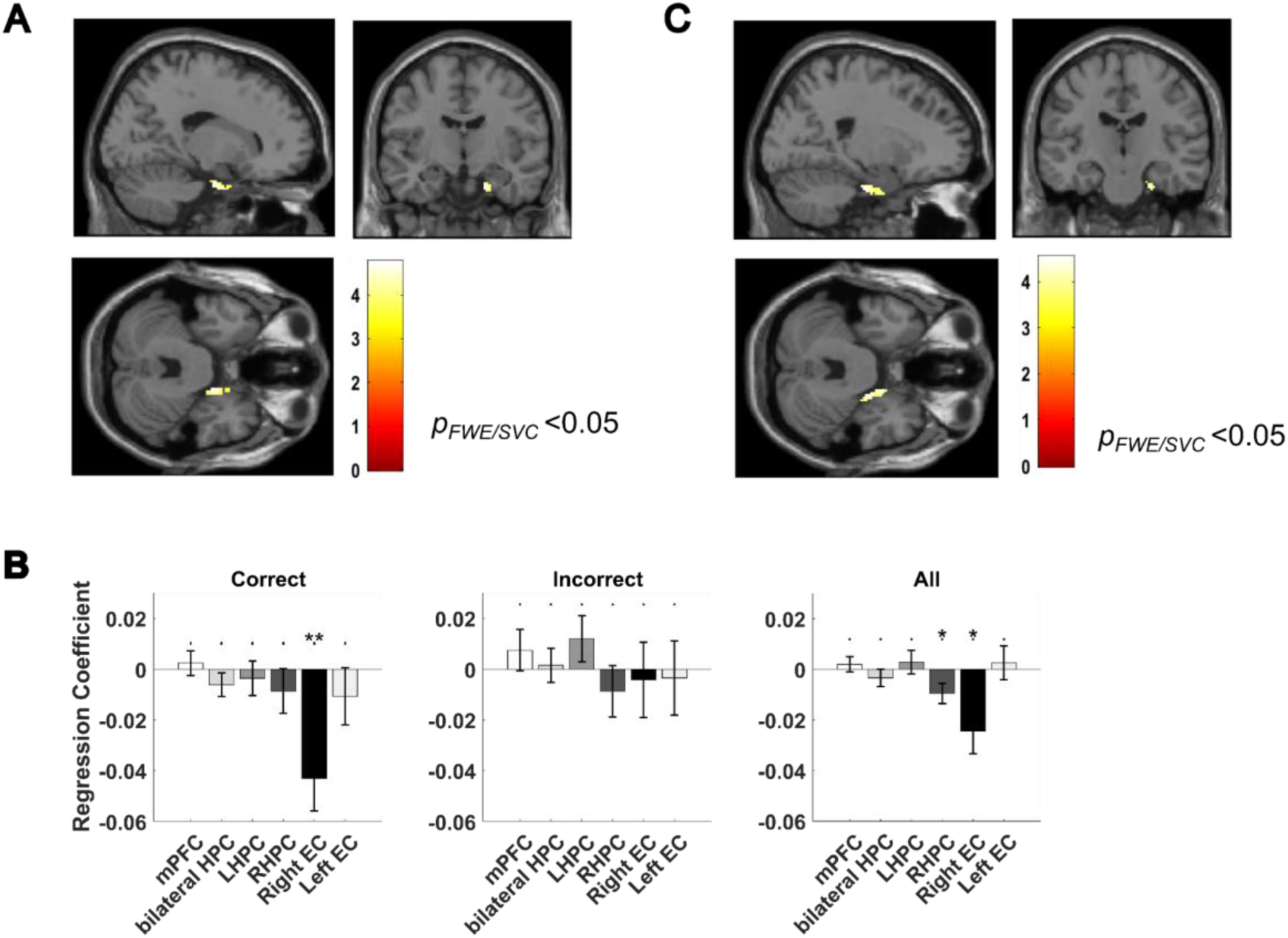
Low theta–fast gamma phase-amplitude coupling (PAC) covaries with distance to the goal during navigation. **(A)** Source localisation of the relationship between 2-5Hz theta - 70-140 Hz gamma PAC and distance to the goal for all correct trials (visualised at *p_uncorr_* < 0.001) within a pre-defined right entorhinal mask. **(B)** Regression coefficients between 2-5Hz theta - 70-140 Hz gamma PAC during navigation and distance to the goal extracted from pre-defined anatomical ROIs across correct trials (left panel), incorrect trials (middle panel), and all trials (right panel). Data shown for the medial prefrontal cortex (mPFC, correct trials: t(22) = 0.490, *p* = 0.629, one-sample t-test; incorrect trials: t(22) = 0.914, *p* = 0.371, one-sample t-test; all trials: t(22) = 0.664, *p* = 0.514, one-sample t-test), bilateral hippocampus (t(22) = −1.31, *p* = 0.203, one-sample t-test; incorrect trials: t(22) = 0.243, *p* = 0.811, one-sample t-test; all trials: t(22) = −0.99, *p* = 0.331, one-sample t-test), right hippocampus (RHPC, t(22) = −0.979, *p* = 0.338, one-sample t-test; incorrect trials: t(22) = −0.85, *p* = 0.403, one-sample t-test; all trials: t(22) = −2.37, *p* = 0.0270, one-sample t-test), left hippocampus (LHPC, correct trials: t(22) = −0.535, *p* = 0.60, one-sample t-test; incorrect trials: t(22) = 1.31, *p* = 0.20, one-sample t-test; all trials: t(22) = 0.611, *p* = 0.548, one-sample t-test), right entorhinal cortex (Right EC, correct trials: t(22) = −3.38, *p* = 0.0027, one-sample t-test; incorrect trials: t(22) = −0.282, *p* = 0.780, one-sample t-test; all trials: t(22) = 2.76, *p* = 0.0115,one-sample t-test) and left entorhinal cortex (Left EC, correct trials: t(22) = −0.95, *p* = 0.355, one-sample t-test, incorrect trials: t922) = −0.226, *p* = 0.824, one-sample t-test; all trials: t(22) = 0.383, *p* = 0.706, one-sample t-test). There is no significant difference between correct trials and incorrect trials in any of the ROIs (*p* > 0.18), although we note that the difference in right entorhinal cortex is marginally significant (t(22) = −2.06, *p =* 0.0578, paired t-test). **(C)** Source localisation of the relationship between 2-5Hz theta - 70-140 Hz gamma PAC and distance to the goal for all correct and novel trials (visualised at *p_uncorr_* < 0.001) within a pre-defined right entorhinal mask.

**Figure S6.**
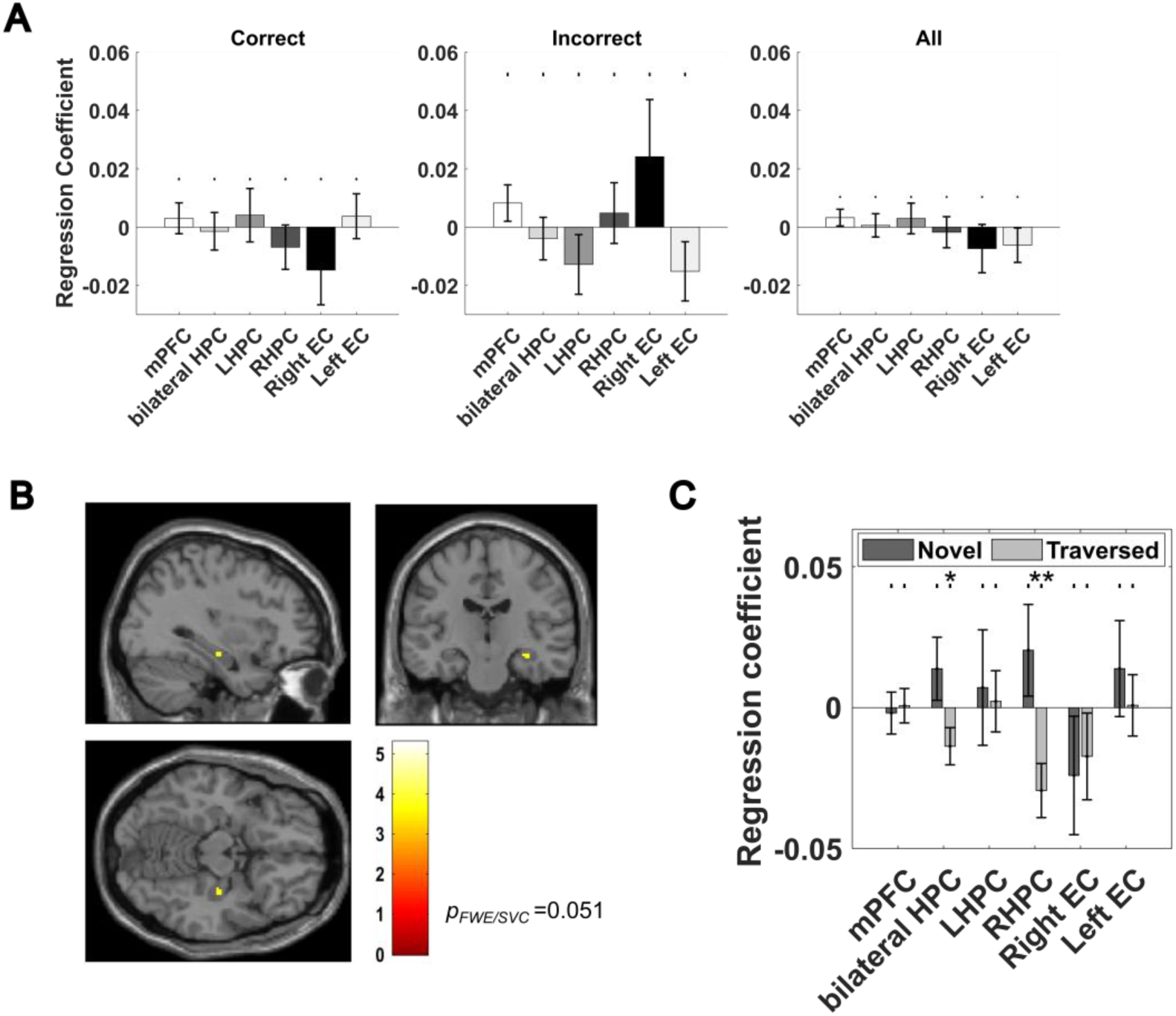
Low theta - slow gamma phase-amplitude coupling covaries with distance to goal during navigation along previously traversed paths. **(A)** Regression coefficients between 2-5Hz theta - 30-70 Hz gamma PAC during navigation and distance to the goal extracted from pre-defined anatomical ROIs across correct trials (left panel), incorrect trials (middle panel), and all trials (right panel). Data shown for the medial prefrontal cortex (mPFC), bilateral hippocampus, right hippocampus (RHPC), left hippocampus (LHPC), right entorhinal cortex (Right EC) and left entorhinal cortex (Left EC). No significant relationship is observed in any ROIs (all *p* > 0.23). **(B)** Source localisation of the relationship between 2-5Hz theta - 30-70 Hz slow gamma PAC and distance to the goal for all correct trials that used previously traversed paths (visualised at *p_uncorr_* < 0.001) within a pre-defined right hippocampal mask. **(C)** Regression coefficient for 2-5Hz theta - 30-70 Hz slow gamma PAC against distance to the goal for correct trials that use novel or previously traversed paths in pre-defined anatomical ROIs. This effect is significantly stronger for previously traversed paths in the right hippocampus (t(22) = −2.26, *p* = 0.0342, paired t-test) and the bilateral hippocampus (t(22) = −2.08, *p* = 0.050, paired t-test), but not in any other ROIs (all *p* > 0.22).

## Notes

### Competing Interest Statement

The authors have declared no competing interest.

